# BIOTIA-DX RESISTANCE Achieved the Best Antimicrobial Resistance Phenotype Prediction Accuracy at CAMDA 2026

**DOI:** 10.64898/2026.05.13.724982

**Authors:** Gábor Fidler, Heather Wells, Ford Combs, John Papciak, Mara Couto-Rodriguez, Sol Rey, Tiara Rivera, Lorenzo Uccellini, Christopher E. Mason, Niamh B. O’Hara, Dorottya Nagy-Szakal, David C. Danko

## Abstract

We present BIOTIA-DX RESISTANCE (BDXR), our submission to the CAMDA 2026 AMR Challenge. This work extends our CAMDA 2025 submission [1] to a new set of six species-drug pairs and adds k-mer-based feature engineering (both targeted and whole-genome) for pairs where the 2025 gene-presence base model underperforms. BDXR achieved a mean accuracy of 86.1% across the six pairs on the CAMDA 2026 test set, ranking first on four pairs, tied for first on *Streptococcus pneumoniae* (penicillin), and second on *Campylobacter jejuni* (nalidixic acid); per-pair test accuracy ranged from 69.9% (*C. jejuni*, nalidixic acid) to 98.8% (*S. pneumoniae*, penicillin). We refer the reader to our 2025 preprint [1] for the underlying workflow, dataset curation, and clinical motivation; this preprint focuses on the results and methodological changes that are new in 2026.

## Background

The biological background of antimicrobial resistance (AMR), the broader motivation for resistance-phenotype prediction, the distinction between marker-gene detection and phenotype prediction, and the overall structure of the CAMDA AMR competition are described in detail in our 2025 preprint [1] and broadly in the literature. Here we summarize only the developments that are specific to CAMDA 2026.

Two relevant developments have occurred since CAMDA 2025. First, the recently released CABBAGE AMR genotype–phenotype database [4] aggregates more than 170,000 sequenced isolates and roughly 1.7M genotype–phenotype pairs across the WHO Bacterial Priority Pathogens, providing the largest public AMR genotype–phenotype resource to date. Second, machine-learning approaches that go beyond classical gene-presence features have begun to appear in CAMDA AMR; notably, Vaska et al. introduced a specialized DNABERT2 large language model fine-tuned with LoRA on de novo bacterial assemblies and achieved 82.8% accuracy and 0.829 macro F1 on the 2025 test set [5].

The CAMDA 2026 AMR competition itself differs from 2025 in two important ways. The species–drug pair set was replaced: 2026 covers *Acinetobacter baumannii* (imipenem), *Klebsiella pneumoniae* (imipenem), *Escherichia coli* (piperacillin/tazobactam), *Campylobacter coli* (tetracycline), *Campylobacter jejuni* (nalidixic acid), and *Streptococcus pneumoniae* (penicillin), spanning gram-positive and gram-negative organisms and both fluoroquinolone- and β-lactam-resistance phenotypes. Models were trained on 800 samples per pair (400 resistant / 400 susceptible) and evaluated on 250 held-out test samples per pair. Unlike 2025, a separate leaderboard was maintained per pair rather than a single overall leaderboard.

### The CAMDA AMR Competition

The Critical Assessment of Massive Data Analysis (CAMDA) is an annual set of competitions for prediction techniques relevant to human health [7]. One of the annual constituent competitions is CAMDA AMR: a competition to predict antimicrobial resistance from microbial genomic data. The CAMDA AMR 2026 competition consisted of predicting resistant phenotype for six clinically relevant species-drug pairs: *Acinetobacter baumannii* (imipenem), *Klebsiella pneumoniae* (imipenem), *Escherichia coli* (piperacillin/tazobactam), *Campylobacter coli* (tetracycline), *Campylobacter jejuni* (nalidixic acid), and *Streptococcus pneumoniae* (penicillin). The set spans gram-positive and gram-negative organisms and includes both fluoroquinolone- and β-lactam-resistance phenotypes. Models were trained on 800 samples per pair (400 resistant / 400 susceptible) and evaluated on 250 held-out test samples per pair. The competition consists of a training set of public data with known resistance phenotype and a test set of newly sequenced data where the resistance phenotype is sequestered.

## Results

### Performance on the CAMDA AMR Training and Test Sets

BIOTIA-DX Resistance achieved a mean accuracy of 86.1% across the six species-drug pairs on the CAMDA 2026 test set, ranking first on four pairs, tied for first on *Streptococcus pneumoniae* (penicillin), and second on *Campylobacter jejuni* (nalidixic acid). Per-pair training and test precision, recall, and accuracy are shown in Table 1.

**Table 1:** Per-pair training and test precision, recall, and accuracy for BDXR on the CAMDA 2026 AMR Challenge. Models were trained on 800 samples per pair (400 resistant / 400 susceptible) and evaluated on 250 held-out test samples per pair. Training scores were obtained by 10-fold GroupKFold cross-validation, with groups defined by mash-distance (sourmash) clusters.

| Species (drug) | Train Precision | Train Recall | Test Precision | Test Recall | Test Accuracy |
| --- | --- | --- | --- | --- | --- |
| *Acinetobacter baumannii* (imipenem) | 0.992 | 0.915 | 0.885 | 0.885 | 0.876 |
| *Klebsiella pneumoniae* (imipenem) | 0.920 | 0.953 | 0.885 | 0.876 | 0.882 |
| *Escherichia coli* (piperacillin/tazobactam) | 0.764 | 0.823 | 0.841 | 0.836 | 0.838 |
| *Campylobacter coli* (tetracycline) | 0.995 | 0.997 | 0.886 | 0.884 | 0.883 |
| *Campylobacter jejuni* (nalidixic acid) | 0.997 | 0.989 | 0.731 | 0.692 | 0.699 |
| *Streptococcus pneumoniae* (penicillin) | 0.960 | 0.960 | 0.988 | 0.988 | 0.988 |
| Overall mean | — | — | — | — | 0.861 |

Test-set accuracy ranged from 0.699 for *Campylobacter jejuni* (nalidixic acid) up to 0.988 for *Streptococcus pneumoniae* (penicillin), with a mean of 0.861. Training scores were obtained via 10-fold GroupKFold cross-validation with mash-distance-based grouping (sourmash), so they do not represent memorization of the training set. As an internal benchmark, our 2025 CAMDA submission used the same gene-presence base model on a different set of nine species-drug pairs and achieved an F1 score of 0.84 on the leaderboard [1]; the 2026 mean test-set accuracy of 86.1% is broadly comparable.

### Performance relative to other submissions to CAMDA AMR

The CAMDA 2026 organizers published a separate leaderboard for each species-drug pair rather than a single overall leaderboard. Table 2 lists each team’s best leaderboard score on each of the six pairs at the time of competition close. Where a team had multiple submissions on a pair, we report their best score. BIOTIA-DX Resistance was top scorer on five of the six pairs (tying with the Yurtseven team on S. pneumoniae and with on C. coli) and was second on C. jejuni (nalidixic acid).

**Table 2:**
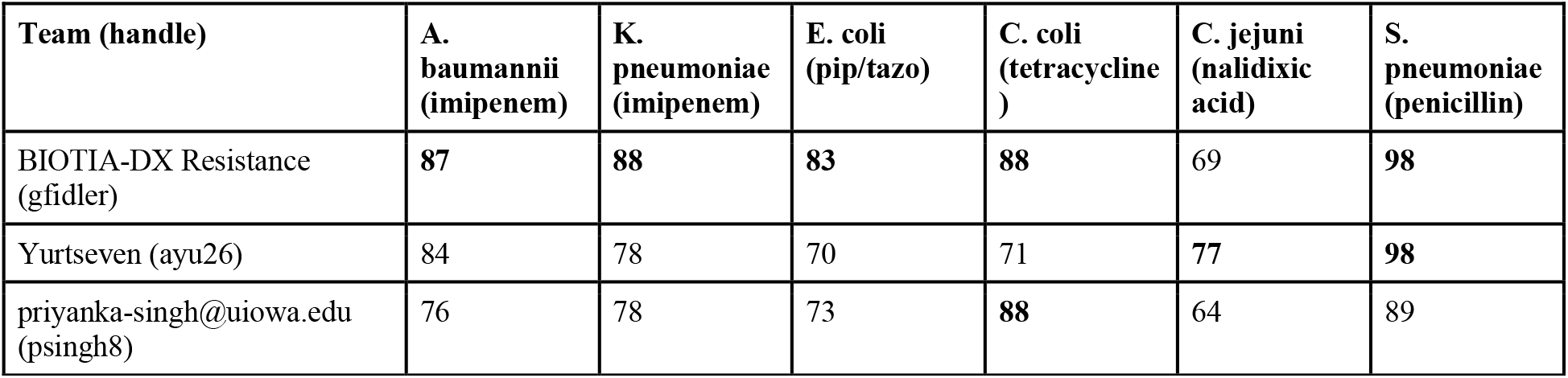
CAMDA 2026 per-pair leaderboard scores by team. The CAMDA 2026 organizers maintained one leaderboard per species-drug pair (no overall score). Each cell shows the listed team’s best score on the corresponding leaderboard at competition close.

## Methods

### Datasets

CAMDA AMR provided two related datasets: a training set and a test set. Both consisted of assembled microbial genomes. The training set contained resistance phenotype given as an ordered categorical variable with *susceptible, intermediate*, and *resistant* values. We treated *intermediate* values as *resistant*. Minimum inhibitory concentration (MIC) values were also given for a subset of samples, but we did not make use of this information.

The test set did only contained genomes, not labels. The resistance phenotypes were known to the CAMDA organizers for each genome but this information was sequestered. The organizers of CAMDA AMR affirmed that phenotypes for the test data did not appear in any public repository. We did not have access to the phenotype data for the test set.

### Training and Evaluation

To train BDXR, we collected over 10,000 genomic sequences with known phenotypes from the supplied CAMDA training data, the Bacterial and Viral Bioinformatics Resource Center (BV-BRC), the Comprehensive Antibiotic Resistance Database (CARD), and other publicly available sources [11], [12]. For the 2026 challenge we additionally incorporated the recently released CABBAGE AMR genotype–phenotype database [4], which aggregates >170,000 isolates and ∼1.7M genotype–phenotype pairs across the WHO Bacterial Priority Pathogens. Data was curated on our platform GeoSeeq. We identified known AMR genes in these genomes and generated clusters of genetically similar genes. Gene clusters were identified based on Jaccard similarity of k-mer spectra. Examples of variants retained on this basis include the *gyrA* Thr86Ile (T86I) substitution underlying nalidixic-acid/fluoroquinolone resistance in *Campylobacter* [8-10], specific *bla*_KPC_ carbapenemase alleles linked to imipenem resistance in *K. pneumoniae* and *A. baumannii*, and altered penicillin-binding-protein alleles (e.g. *pbp1a*) for penicillin-non-susceptible *S. pneumoniae*.

Each genome was described as a vector of gene presence or absence. For gene families where we determined variants to be relevant, we selected variants to include in the vector by filtering to only those which were statistically significantly associated with the resistant phenotype. Vectors were subject to unsupervised dimensionality reduction. The resulting vectors were paired with susceptibility data as labels and used to develop predictive models.

Using the input vectors described above, we trained a family of machine learning models for taxonomic and drug resistance groups. Model input was filtered based on feature importance to reduce dimensionality and training was subject to 10-fold GroupKFold cross-validation, with groups defined by mash-distance (sourmash) clusters so that splits remain phylogeny-aware, to reduce overfitting. Once features were selected, models were subject to hyperparameter tuning via GridSearchCV and an optimal prediction probability threshold was selected based on F1 score. Model selection was performed by training multiple models and selecting the best performer; for 2026 the candidate set comprised our base gene-presence model and two new k-mer-based variants (targeted and whole-genome), with the final per-pair model chosen on cross-validated F1. Training scores were obtained by averaging 10-fold GroupKFold cross-validation data.

### Data Cleaning

All genomic data was rigorously cleaned before processing. Raw input reads from public data were filtered to remove low-quality reads, repetitive reads, and reads that mapped to the human reference genome. Remaining reads were assembled into contigs and putative genes were called. The CAMDA competition provided pre-assembled genomes for its training and test data so these steps were omitted from our workflow for those data. Genes were compared to known AMR clusters to identify similar but possibly undescribed AMR genes.

After training we generated phenotype predictions for the CAMDA test set. These predictions were submitted to the CAMDA leaderboard which automatically generated an overall score. Competition rules allowed up to three submissions to the test set and we submitted twice.

## Discussion

Resistance prediction from genetic data remains a moving target. While accuracy has improved across successive CAMDA AMR challenges, many species/drug pairs are not consistently at clinical-grade accuracy on a held-out test set. Continued progress will depend on larger and more carefully curated genotype–phenotype datasets, more robust evaluation protocols, and steady methodological improvement. The cumulative trajectory across years of CAMDA AMR suggests that incremental gains, sustained by multiple groups, are the realistic mode of progress in this area.

Deep-learning-based approaches, both AMR-specific models such as DNABERT2 [5] and broader genome foundation models such as GenomeOcean and Evo 2 [13,14], are increasingly active in CAMDA and elsewhere, and we expect them to play an increasingly central role in future submissions. Our group is developing deep-learning approaches in parallel with the workflow described here. As discussed in [1], mechanistic studies of AMR are likely to remain important alongside such models, since they are largely non-explanatory and unlikely to fully substitute for biological understanding.

## Data Availability

The CAMDA AMR Leaderboards are publicly available on the CAMDA website at https://bipress.boku.ac.at/camda2026/competitions/aba_imi/?leaderboard https://bipress.boku.ac.at/camda2026/competitions/cco_tet_2026/?leaderboard https://bipress.boku.ac.at/camda2026/competitions/cje_na_2026/?leaderboard https://bipress.boku.ac.at/camda2026/competitions/ecopip_2026/?leaderboard https://bipress.boku.ac.at/camda2026/competitions/kpnimi_2026/?leaderboard https://bipress.boku.ac.at/camda2026/competitions/spn_pen_2026/?leaderboard

